# Probing the Evolution of Genes Associated With DNA Methylation in *Listeria monocytogenes*

**DOI:** 10.1101/2023.11.04.565605

**Authors:** Ogueri Nwaiwu, Helen Onyeaka, Catherine Rees

## Abstract

In the last decade, there have been increased reports of atypical *Listeria* and the discovery of new species. There are public health concerns that new strains may come with increased pathogenicity. Hence, this study aimed to establish the prevalence, evolutionary lineage and ancestry of a *Listeria monocytogenes* collection that includes isolates that harbour a unique set of methylase genes. The addition of methyl groups to DNA can interfere with transcription. Allelic-specific lineage analysis and ribotyping with southern hybridization were carried out after which further phylogenetic analysis was performed *in silico*. Results show that all the methylase strains belonged to Lineage I and were serotypes 4b or 4d. All designated ancestral strains also belonged to Lineage 1. A *Listeria monocytogenes* plasmid from a serotype 1/2a (Lineage II) contained sequences homologous to that of Lineage I isolates. The methylase nucleotide sequence in the strains studied appears to be highly conserved in *Listeria monocytogenes* and not yet orthologous among other bacterial genera. It is of epidemiological interest and public benefit if wider or continuous surveillance is carried out to ascertain if these rare strains are linked with increased pathogenesis, food type or geographical region.

## 1. Introduction

*L. monocytogenes* remains an important foodborne pathogen that contaminates food at any stage during production, processing, and storage, resulting in potential food safety issues. This enables its detection in food, environmental and clinical samples [1]. The organism is among the most common causes of foodborne disease in humans, causing death and hospitalization with high costs, making rapid detection important [2]. This bacterium serves as a valuable microorganism model for investigating the interplay between a pathogen and its host due to its unique ability to colonize the intestinal lumen, cross various barriers such as the intestinal, blood-brain, and placental, and cause bacteremia [3]. The ability to survive adverse conditions is one of the main reasons *L. monocytogenes* persists in the food-processing environment. The microbe is linked with being baro-tolerant [4], and survives under low temperatures and pH. Also, it is believed to have a dual intercellular lifestyle, which helps it to invade, colonise and establish in a host [5]. Common food that can easily be contaminated includes dairy [6], vegetables [7, 8], meat and poultry [9,10]. A recent survey [11] found that serogroup 1/2a-3a dominate food, 4a-4c in livestock, and 1/2a-3a in humans. Also, the dominance of *L. monocytogenes* lineage II in food, humans, and lineage III in livestock was observed.

Contamination of food by *L. monocytogenes* is possible because it can survive stress in the food processing environment. It has been reported that one of its control mechanisms that ensure its survival in processing stress is through the production of heat and cold-shock proteins [12], and exopolymeric substances, which hold its biofilm firmly to a surface [13]. It is known that once persistence has been established, the organism will continue to survive cleaning and sanitation over a period [14]. This shows the importance of monitoring prevalence to enable informed public health decisions by policymakers and clinicians [15]. A gold standard approach for monitoring different *L. monocytogenes* in the food chain is using whole genome sequencing [16], which has enabled scientists to gain insights into the fundamental characteristics of different *L. monocytogenes* isolates. The study of genetic profiles of bacteria, in general, has revealed that DNA can be affected by environmental factors without the sequence being affected. This is commonly known as epigenetics good example is DNA methylation, a process that allows the addition of methyl groups to DNA.

As Rajeev et al. [17] pointed out, DNA methylation is a reversible epigenetic modification of DNA and is associated with the dynamic regulation of gene expression. In bacteria [18] and mammals [19], adenine and cytosine are known to be methylated. Even in plants, a report [20] highlighted that DNA methylation can be used as a marker to monitor genome evolution and stress adaptation to aid crop disease resistance and improvement. Modifying DNA with a methyl group, which is catalysed by DNA methyl transferase is one of the most occurring nucleotide modifications present in genomes of both prokaryotes and eukaryotes [21]. Also, it is essential for normal development and long-term transcriptional silencing [22], and understanding human DNA methylation patterns helps analyse the molecular mechanism of diseases[23,24]. It has been reported [25] that in bacteria genomes, C5-methyl-cytosine, N4-methyl-cytosine, and N6-methyl-adenine can be formed after replication and that these base methylation can modulate the interaction of DNA-binding proteins with their cognate sites which leads to the formation of epigenetic lineages by phase variation. Others [26–28] have elucidated further phenotypic and epigenetic regulation importance of methylation in bacteria.

In *L. monocytogenes* patho-epigenetics, different strategies are used to control the expression of host genes during invasion [29]. These include activating cytosolic signaling pathways at the chromatin and targeting epicactors in the nucleus. The host can respond to *Listeria* invasion by undergoing transcriptional changes and activating protective genes [30]. An investigation [31] has revealed diversity in the restriction-modification systems and DNA methylation sites in a study of 15 *L. monocytogenes* genomes. Importantly, it was found that some strains exhibited no methylation, even though they harboured methyltransferase genes. Historically, although the occurrence of 5-methylcytosine was first reported [32] several decades ago in bacteria, the prevalence, occurrence and role in *L. monocytogenes* require further studies. This is very important, considering that the role of methylation in *L. monocytogenes* may vary among its lineages.

In the last decade, the number of *L. monocytogenes* has increased [33], with the 27th species, *Listeria Ilorinensis* reported [34] as of June 2022. The incidence of *L. monocytogenes* with atypical features is also on the rise, hence continuous surveillance and monitoring are important for any pathogenic link with new methylase strains as they evolve. In a previous study, a cytosine methyltransferase present in *L. monocytogenes* was identified and the nucleotide sequence that encodes the lcmA gene was deposited (AJ302030.1) [35]. In this study, our aim was to determine the prevalence and evolutionary history of these genes in a collection of L. monocytogenes strains.. Following confirmation of the lineage of the strains, the possibility of the strains under study being the rare serotype 4b Lineage III strains was explored, and the most probable evolutionary ancestors were pointed out. The ancestral strains, clonal complex and multilocus sequence types of strains that harbour the nucleotide sequences studied and their pathogenic potential were highlighted after further *in silico* analysis.

## 2. Results

### 2.1. Strains with methylase genes

Using primer sequences in Table 1, strains under study were screened to ascertain prevalence. Amplification of the methylase genes was observed only in some strains of PCR serotype 4b and was negative for all isolates of serotypes 1/2a or 1/2b. The methylase genes were detected in strains from food (Lm13), veterinary (Lm17), and the clinical strains isolates Lm 23, 24 and 25 (Table 1). The well-known isolates in literature (Lm 10403S and Lm EGD; 1/2a) and the reference strain ATCC 23074 (4b) did not amplify the primers used. Overall, only five out of the 45 (11%) strains screened had the genes in their chromosome. A representative electrophoresis output showing the methylase gene fragments amplified at approximately 1000kb is shown in Figure 1.

**Table 1.** Methylase gene amplification and lineage classification of *Listeria monocytogenes* strains from various sources.

| Strain | Molecular Serotype | Source | Methylase | Lineage Classification |  |  |  |  |  |  | Lineage |
| --- | --- | --- | --- | --- | --- | --- | --- | --- | --- | --- | --- |
|  |  |  |  | *actA1* | *plcB2* | *actA3* | *ORF 2110* | *lmo1134* | *lmo 2821* | *lmo 0733* |  |
| Lm1 | 1/2a | Milk | Negative | - | + | - | nd | nd | nd | nd | II |
| Lm2 | 1/2a | Milk | Negative | - | + | - | nd | nd | nd | nd | II |
| Lm3 | 1/2a | Milk | Negative | - | + | - | nd | nd | nd | nd | II |
| Lm4 | 1/2b | Milk | Negative | + | - | - | nd | nd | nd | nd | I |
| Lm5 | 1/2a | Milk | Negative | - | + | - | nd | nd | nd | nd | II |
| Lm6 | 4b | Milk | Negative | + | - | - | + | + | + | + | I |
| Lm7 | 4b | Milk | Negative | + | - | - | + | + | + | + | I |
| Lm8 | 4b | Milk | Negative | + | - | - | + | + | + | + | I |
| Lm9 | 1/2a | Food | Negative | - | + | - | nd | nd | nd | nd | II |
| Lm10 | 1/2a | Food | Negative | - | + | - | nd | nd | nd | nd | II |
| Lm11 | 1/2a | Food | Negative | - | + | - | nd | nd | nd | nd | II |
| Lm12 | 1/2a | Food | Negative | - | + | - | nd | nd | nd | nd | II |
| Lm13 | 4b | Food | Positive | + | - | - | + | + | + | + | I |
| Lm14 | 4b | Food | Negative | + | - | - | + | + | + | + | I |
| Lm16 | 1/2a | Veterinary | Negative | - | + | - | nd | nd | nd | nd | II |
| Lm17 | 4b | Veterinary | Positive | + | - | - | + | + | + | + | I |
| Lm18 | 1/2a | Veterinary | Negative | - | + | - | nd | nd | nd | nd | II |
| Lm19 | 1/2a | Veterinary | Negative | - | + | - | nd | nd | nd | nd | II |
| Lm20 | 1/2b | Clinical | Negative | + | - | - | nd | nd | nd | nd | I |
| Lm21 | 1/2a | Clinical | Negative | - | + | - | nd | nd | nd | nd | II |
| ATCC-23074 | 4b | Clinical | Negative | + | - | - | + | + | + | + | I |
| Lm22 | 1/2c | Clinical | Negative | - | + | - | nd | nd | nd | nd | II |
| Lm23 | 4b | Clinical | Positive | + | - | - | + | + | + | + | I |

**Table 1.** *Contd.*
| Strain | Molecular Serotype | Source | Lineage Classification |  |  |  |  |  |  |  |  |
| --- | --- | --- | --- | --- | --- | --- | --- | --- | --- | --- | --- |
|  |  |  | *meth* | *actA1* | *plcB2* | *actA3* | *ORF 2110* | *lmo1134* | *lmo 2821* | *lmo 0733* | Lineage |
| Lm24 | 4b | Clinical | Positive | + | - | - | + | + | + | + | I |
| Lm25 | 4b | Clinical | Positive | + | - | - | + | + | + | + | I |
| Lm26 | 1/2a | Food | Negative | - | + | - | nd | nd | nd | nd | II |
| Lm27 | 1/2c | Food Env | Negative | - | + | - | nd | nd | nd | nd | II |
| Lm28 | 1/2a | Food | Negative | - | + | - | nd | nd | nd | nd | II |
| Lm29 | 1/2a | Food | Negative | - | + | - | nd | nd | nd | nd | II |
| Lm30 | 1/2a | Food | Negative | - | + | - | nd | nd | nd | nd | II |
| LO28 | 1/2c | - | Negative | - | + | - | nd | nd | nd | nd | II |
| Lm EGD | 1/2a | - | Negative | - | + | - | - | + | + | + | II |
| Lm10403S | 1/2a | Clinical | Negative | - | + | - | nd | nd | nd | nd | II |
| Lm00101-0305 | 4b* | Food Env | Negative | + | - | - | + | + | + | + | I |
| Lm00048-0305 | 1/2c* | Food Env | Negative | - | + | - | nd | nd | nd | nd | II |
| Lm0054-0305 | 1/2b* | Food Env | Negative | + | - | - | nd | nd | nd | nd | I |
| Lm00064-0305 | 4b* | Food Env | Negative | + | - | - | + | + | + | + | I |
| Lm000630-0305 | 1/2c* | Food Env | Negative | - | + | - | nd | nd | nd | nd | II |
| Lm000622-0305 | 1/2a* | Food Env | Negative | - | + | - | nd | nd | nd | nd | II |
| Lm000636-0305 | 1/2c* | Bakery | Negative | - | + | - | nd | nd | nd | nd | II |
| Lm0006-0305 | 1/2c* | Bakery | Negative | - | + | - | nd | nd | nd | nd | II |
| Lm00006-0305 | 1/2c* | Bakery | Negative | - | + | - | nd | nd | nd | nd | II |
| LmNG1 | 1/2b** | Vegetable | Negative | + | - | - | nd | nd | nd | nd | I |
| LmNG2 | 1/2b** | Vegetable | Negative | + | - | - | nd | nd | nd | nd | I |
| LmNG3 | 1/2a** | Vegetable | Negative | - | + | - | nd | nd | nd | nd | I |

nd = not determined; Env = environment; \*Molecular serotype reported in a previous study [7]
\*\* Molecular serotype and lineage reported in another report [36].

**Figure 1.**
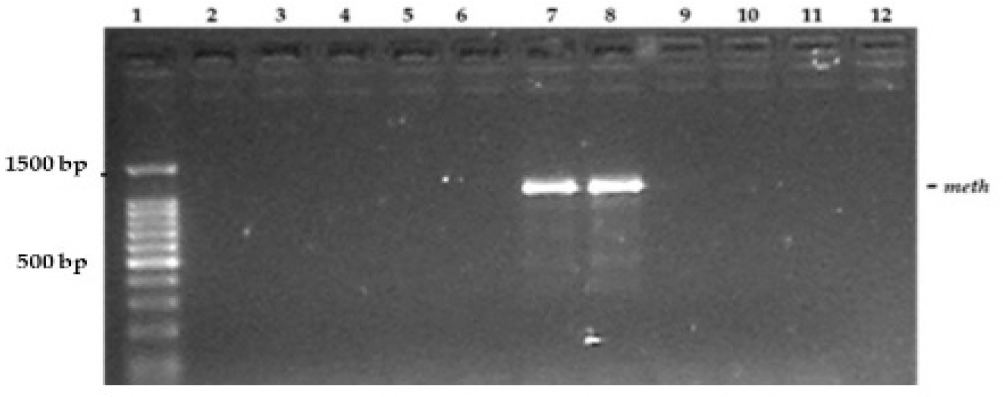
Screening of strains for genes associated with DNA methylation. Lane1=100 bp marker; 2=Lm GD(1/2a), 3=Lm10403S(1/2a), 4=Lm0006(1/2c), 5=Lm000101(4b), 6=Lm00006(1/2c), 7=Lm13(4b), 8=Lm 24(4b), 9= Lm00064(4b), 10= Lm23074(4b), 11=Lm 000622(1/2a), 12= No DNA template. 5μl of PCR product was loaded with 1μl of dye and expressed in 2% agarose gel.

### 2.2. Lineage classification of strains

Lineage classification was carried out by looking at specific alleles. The evolutionary lineage classification outcome for strains using allelic-specific oligonucleotide PCR analysis is shown in Table 1. It was found that all strains that are serotypes 1/2b and 4b amplified at 373bp expected for lineage I, while strains that are serotypes 1/2a and 1/2c amplified at 564bp expected for lineage II. There was no detection of the rare lineage III specific sequence since the gene *actA3* (277kb) was not amplified in any of the samples analysed in this study (Figure 2). Further lineage analysis carried out to differentiate between lineage 1 and III serotypes strains showed that all the serotype 4b strains analysed including the isolates harbouring the methylase genes sequence amplified the genes *ORF 2110, lmo1134, lmo2821* and *lmo 0733* (Table 1, Figure 3A-D), which confirmed that they were lineage I isolates. There were no *ORF 2110* fragments in the reference strain Lm EGD (1/2a) used as a control (Figure 3a).

**Figure 2.**
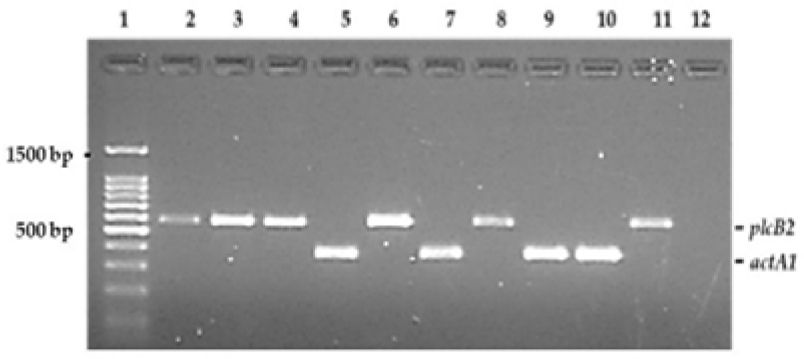
Lineage classification of *L. monocytogenes* by allelic specific oligonucleotide PCR (ASO-PCR) of the virulence gene clusters of prfA. Lanes 1=1kb marker, 2=LmEGD (1/2a), 3=Lm10403S (1/2a), 4=Lm0006 (1/2a), 5=Lm13 (4b), 6=Lm21 (1/2a), 7=Lm23 (4b), 8=Lm00048 (1/2c), 9=Lm74, 10=Lm000101 (4b), 11=Lm000622 (1/2a), 12=negative control. 5μl of the digest was added to 1μl of loading dye before loading into each well. The PCR products were expressed on 1.5% agarose gel and scored against a 100 bp ladder.

**Figure 3.**
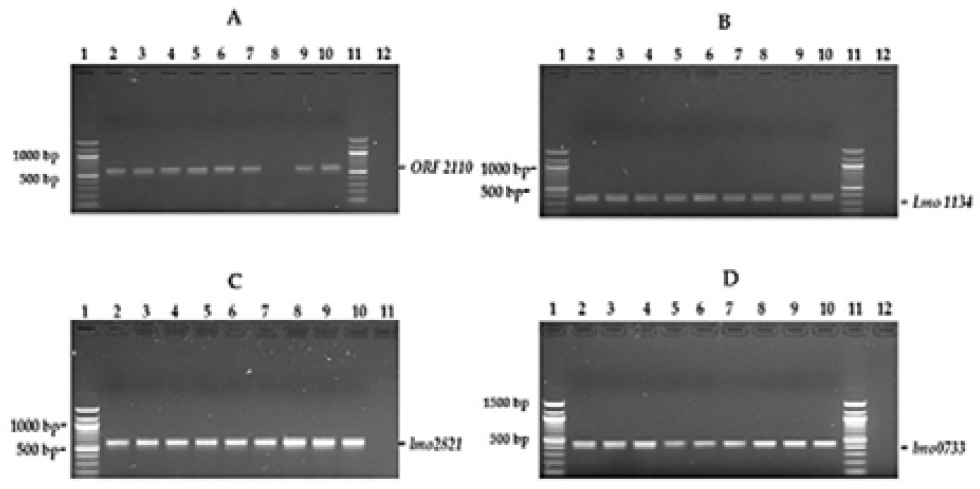
Checking serotype 4b strains for rare Lineage III fragments. In all panels above -Lane1=100 bp marker; 2=Lm 13 (4b), 3=Lm17 (4b), 4=Lm23 (4b), 5=Lm24 (4b), 6=Lm25 (4b), 7=Lm23074, 8=Lm EGD (1/2a), 9= Lm000101 (4b), 10= Lm00064 (4b), 11=100 bp marker, Lane 12= No DNA template. Targeted genes were *ORF* 2110 **(A)**, *lmo1134* (B) *lmo2821* (C) and (D) *lmo0733*.

### 2.3. Ribotype profile of strains harbouring the methylase genes

Ribotyping by Southern hybridization was carried out to establish if the methylase strains were genetically from the same clone. After chromosomal DNA was digested with *Eco*RI, it was transferred to a nylon membrane and then hybridized with a 16S rRNA probe, different profiles emerged for strains analysed. The serotype 4b strains, which contain the methylase genes, had the same profile (Lanes 5, 6, 7, Fig. 4). The profiles of the 4b serotypes were different to that of serotypes 1/2c or 1/2a.

**Figure 4.**
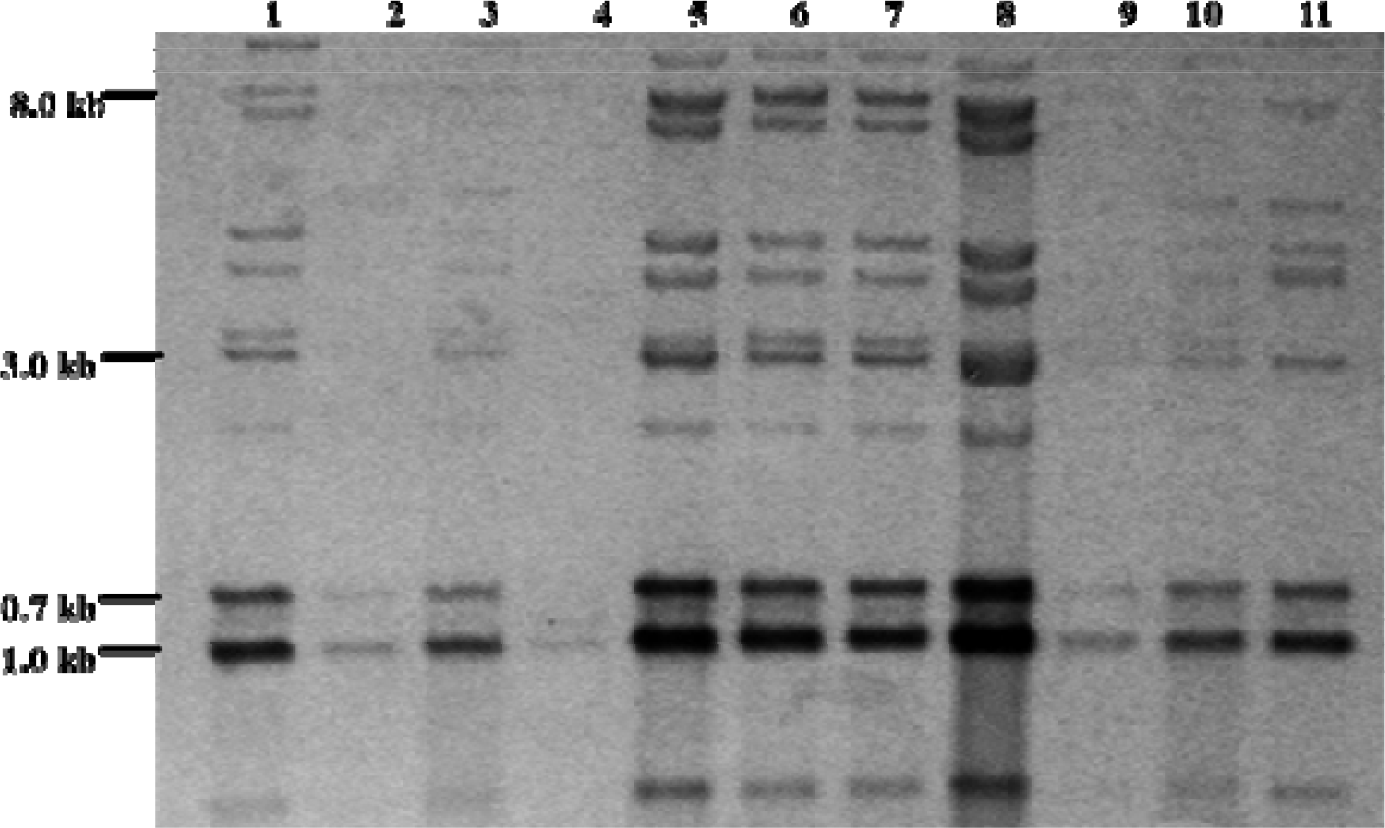
Figure 3.5.1 Southern hybridization of EcoRI digested chromosomal DNA electrophoresed in 0.7% agarose gel, transferred to a nylon membrane and hybridized with DIG labelled 16S RNA probe. The temperature of hybridization was 4 2°C. Molecular weight sizes in Kb are shown on the left of the figure. 1=Lm13 (4b), 2=Lm27 (1/2c), 3=Lm000101 (4b), 4=Lm0006 (1/2c), 5=Lm17 (4b), 6=Lm23 (4b), 7=Lm24 (4b), 8=Lm4 (1/2b), 9=Lm000622 (1/2a), 10=Lo28 (1/2c), 11=Lm00064 (4b). The profiles of methylase 4b strains (lanes 5, 6, 7) are different to that of another 4b strain (Lane 11).

**Figure 5.**
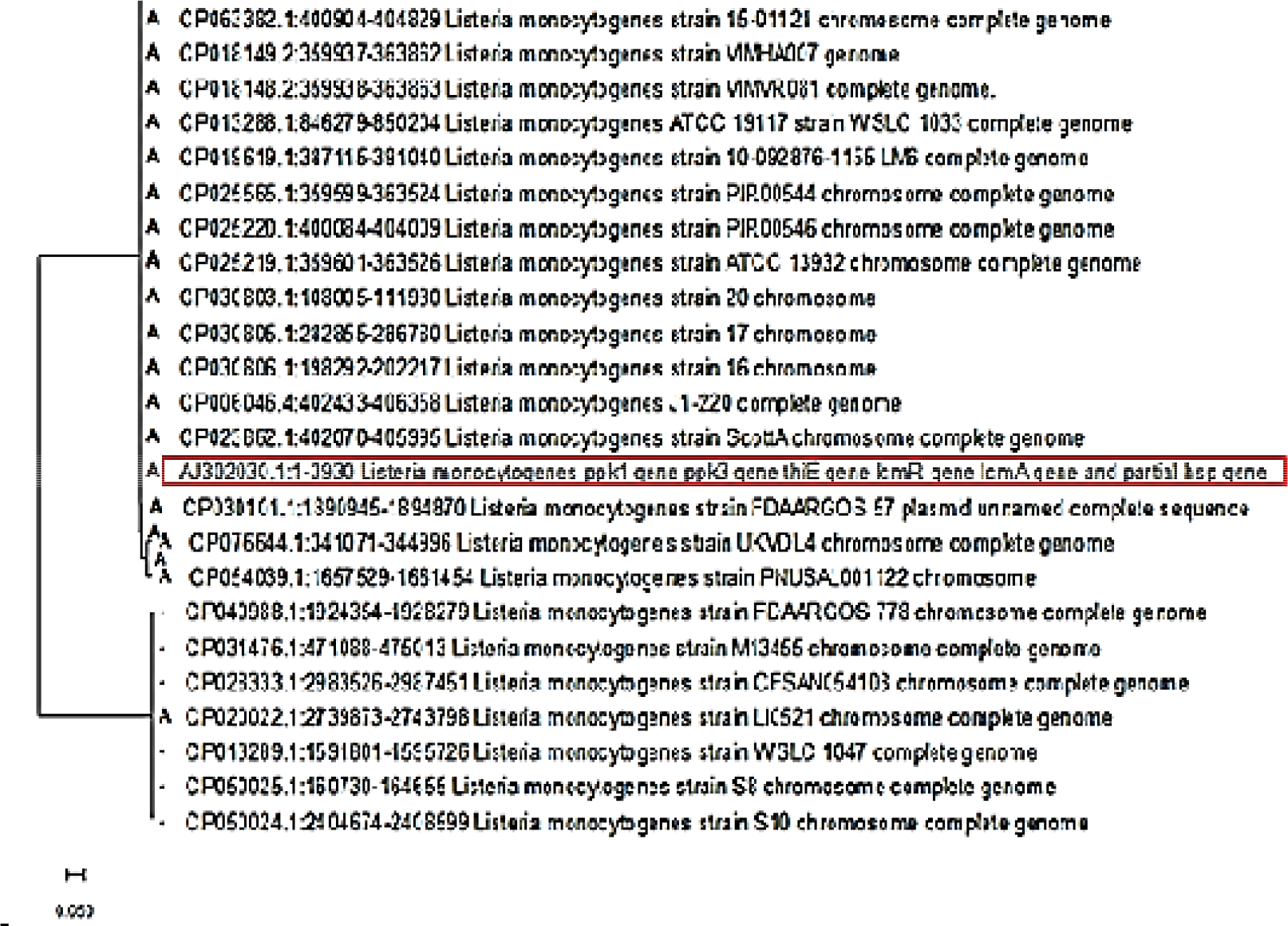
Phylogenetic tree showing the evolutionary history of *L. monocytogenes* strains containing the nucleotide sequence of genes (in red box) associated wit cytosine DNA methylation. The tree was constructed after alignment was built with MUSCLE and a test of phylogeny was carried out using the maximum likel hood statistical method based on the Tamura-Nei model. The bootstrap iteration was set at 1000 iterations. The tree is drawn to scale, and the analysis was carried o with 24 nucleotide sequences. Strains with ‘‘A’’ before them were designated as ancestors of the reference sequence (in red box) following analysis with the ancestr function of MEGA 11 computer software.

### 2.4. Prevalence and probable ancestors of strains harbouring the methylase genes

To further ascertain the global prevalence of strains harbouring the complete nucleotide sequence of genes associated with DNA methylation (GenBank-number AJ302030.1) in *L. monocytogenes*, a search was carried out in the NCBI BLAST ® suite. The outcome of the search (Table S1) showed that sequence similarity or per cent identity was low in other genera of bacteria like *Escherichia, Enterococcus, Klebsiella, Serratia, Lactococcus*, and *Salmonella*. Also, it was low in other *Listeria* species like *L. welshimeri, L. innocua* and *L. marthi* (≤ 93%). Interestingly, strains of *Pseudoalteromonas carrageenovora* had more similarity (96%) than some *L. monocytogenes* serotypes 1/2c or 1/2a. A narrow pool of *L*.monocytogenes had high similarity (99%), mainly serotype 4b isolates and a couple of 4d strains. The high-scoring isolates were selected, and their ancestral relationships and history was searched using MEGA (version 11) software. Following the construction of a phylogenetic tree (Fig.5), it was found that two distinct clades were formed. All the strains that fell under the same clade with the sequence of interest were designated ancestors after *in silico* analysis. Other strains, which were not designated as an ancestor, fell into the other clade with just one strain that had ancestral features.

Further searches on the *Listeria* PasteurMLST database for other evolutionary features of the aforementioned isolates were carried out. Ancestral strains under study with records available in the database, all belonged to clonal complex 2 and epidemiological clone IV. However, their MLST and cgMLST profiles differed (Table 2).

**Table 2.** Clonal complex, sequence type, and epidemic clone of some strains containing genes. associated with DNA methylation.

|  | *L. monocytogenes* strains | % Identity | Accession | Serotype | PCR-se<br>rotype | Clonal<br>complex | cgmlst | MLS<br>T<br>(ST) | Source | Epidemi<br>c clone | Reference |
| --- | --- | --- | --- | --- | --- | --- | --- | --- | --- | --- | --- |
| **1** | ScottA chromosome,<br>complete genome | 99.8 | CP023862.<br>1 | 4b | IVb | 2 | 416 | 290 | Unknown | IV | [37] |
| **2** | WSLC 1047 chromosome,<br>complete genome<br>(non-virulent) | 99.8 | CP013289.<br>1 | 4d | IVb | 2 | 375 | 290 | Unknown | - | [38] |
| **3** | S8 chromosome, complete<br>genome | 99.8 | CP050025.<br>1 | 4b | IVb | 2 | 4059 | 2 | Envirenment<br>al floor | - | [39] |
| **4** | VIMVR081, complete<br>genome | 99.8 | CP018148.<br>2 | 4b | IVb | 2 | - | 145 | Rodent | - | [40] |
| **5** | VIMHA007 chromosome | 99.8 | CP018149.<br>2 | 4b | IVb | 2 | - | 2 | Stillborn | - | [40] |
| **6** | ATCC 19117, serotype 4d | 99.8 | FR733643.<br>1 | 4d | IVd | 2 | 405 | 2 | Sheep | - | [41] |
| **7** | 15-01121 chromosome,<br>complete genome | 99.8 | CP063382.<br>1 | 4b | IVb |  |  | 2 | Clinical | - | [42] |
| **8** | J1-220 complete genome | 99.8 | CP006046.<br>4 | 4b | IV | 2 | - | - | Vegetables | IV | [43] |
| **9** | LI0521 complete genome | 99.8 | CP020022.<br>1 | 4b |  | 2 | - | - | Clinical-cont<br>aminated<br>cheese | IV | [44] |

## 3. Discussion

In order to identify which strains in our *L. monocytogenes* collection carried the methylase genes, we utilized the methylase primers outlined in Table 4 to analyze the genomic DNA from all strains being studied. Our results indicated that the sequence of interest was exclusively present in strains of serotype 4b, providing further evidence that the methylase genes under investigation were predominantly present in this group.. As methylation is an epigenetic event, the trigger in one strain may differ from another. The result is similar to that of others [31] in that their investigation found a high diversity in methylation statuses despite high degrees of genome conservation. The authors investigated 15 genomes and found that eleven strains possessed a single methyltransferase, and one strain contained two methylation systems. In addition, three isolates showed no methylation even though they had methyltransferase genes and in three isolates, unknown novel methylation patterns were observed. The report concluded that the epigenetic DNA methylation sites in *L. monocytogenes* are very diverse. A study of other *L. monocytogenes* collections around the globe may be beneficial to ascertain the real prevalence and occurrence of similar methylase probes used in our investigation. Though many studies have been conducted on DNA methylation in bacteria, the mechanistic knowledge underpinning the process is limited [45]. Others have proffered the explanation for some epigenetic mechanisms that may occur in bacteria and possibly cause methylation in host-pathogen relationships. These include diet and gut microbiota [46-48].

To ascertain if the strains that possess the methylase gene belonged to the same evolutionary lineage, allelic-specific analysis was carried out, after which strains of serotype 4b were probed to establish if any of them belong to lineage III rather than I. The result, which showed that all the 4b strains analysed in this study (both methylase positive and negative isolate) belong to lineage I, is consistent with earlier findings of Ward et al. [49] and Orsi et al. [50] that placed 4b isolates under lineage 1. Another report [51] highlights that Lineage I isolates includes major epidemic clones of *L. monocytogenes* associated with human listeriosis cases. Lineage III isolates were not detected in this study, possibly because they occur mainly in serotypes 4a and 4c as shown by others [52]. Following an analysis [53] of the four lineages of *L. monocytogenes*, it has been established that there are differences in the recombination rate, and the core genome variation between the lineages exhibited a distinct pattern. A review [54] highlighted that phase variation may distort methylation target specificity and cause bacteria to switch between alternative DNA methylation profiles. Another report [55] posits that in the host genome, methyl groups could be added through type I restriction-modification systems, which can add methyl groups to the host genome. The characteristics of *L. monocytogenes* lineages and serotypes in different environmental conditions are reported widely in several studies [56-61]

Since the methylase-positive strains were not found to be the rare lineage III serotype, we carried out ribotyping with Southern hybridization. The profile obtained was the same for the methylase strains analysed, which supported the hypothesis that the isolates were from the exact epidemiological clone. However, Casadesús and Low [62] posited that even though bacterial populations are clonal, the formation of distinct bacterial lineages may appear during harsh conditions and that inheritable phenotypic diversity aided by perpetuating methylation patterns may occur without any alteration of the DNA sequence. The limitation of this assay was that clonality evidence was based on five methylase-positive strains, which were available for testing and hence more evidence was required. To this end, a search was carried out on the NCBI database to check for the prevalence of the nucleotide unit (AJ302030.1) containing the characterized methyl transferase [35]. Although, the search was optimized for 5000 isolates, only 396 strains contained some or full segments of the query sequence. Just 25 strains had 100% homology or query coverage (query length included in the aligned segments) and up to 99% identity (Table S1), which suggests that the sequence of interest was highly conserved in *L. monocytogenes*. This evidence is supported by the fact that other *Listeria* species (Welshimeri and Marthii, which had 100% homology had poor per cent sequence identity (93%). In addition, no other bacteria genera had any significant homology or per cent identity (Table S1). When the documented serological features were examined, it was found that strains other than serotype 4b also had 100% homology and 99% identity. This included strains WSLC 1047 and ATCC 19117, which are nonvirulent *L. monocytogenes* strains of serotype 4d. This is not too surprising considering that serotypes 4b and 4d are in the same lineage I as reported by Rawool et al. [52]. However, the main highlight of the *in silico* analysis is that an unnamed plasmid belonging to *L. monocytogenes* FDAARGOS_57 serotype 1/2a which is in lineage II had the methylase sequence albeit at a slightly lower per cent identity (98.7%; S1) but with 100% sequence similarity. It is possible the plasmid gained the sequence through horizontal gene transfer. Plasmids in the environment are notorious for acquiring antimi-crobial-resistant genes through horizontal gene transfer.

Further phylogenetic analyses carried out with MEGA (Version 11) on the strains with high sequence similarity (100%) and no less than 98% identity, showed that the plasmid was as an ancestor and fell in the same clade with most of the serotype 4b strains designated as ancestors. Interestingly some 4b strains that were in another clade were not designated as ancestors even though they possess the methylase sequence. This indicates that the methylase strains are beginning to acquire distinct features, which may be due to environmental pressure. Further work may establish if this is due to horizontal gene transfer.

To further explore the properties of the *L. monocytogenes* isolates with the methylase sequence, historical records in the *Listeria* database [63] were consulted. The database currently has up to 2049 genomes (as of 21.12.22). This effort was limited because only a few of the strains used in constructing the phylogenetic tree in this study had records. Nevertheless, a clear pattern emerged in that all the strains found on the database were from clonal complex 2 [64], which supports the similar hybridisation profiles in methylase strains. However, four different cgMLST types (375, 405, 416, 4059) and three STs (2, 145, 290) were recorded for some of the strains. It has been noted [65] that strains may be in the same clone and have very similar genetic backgrounds, but they often cannot be distinguished using various molecular subtyping methods.

This heterogeneity observed in strains from the same lineage is consistent with the findings of Moura et al. [66]. The authors demonstrated heterogeneity and other varying genomic features of strains from different environments. Furthermore, an analysis [65] of 4b serotypes and their variants (4bV) concluded that 4bV strains have undergone adaptive responses to selective pressures that could help their survival in the environment while retaining the pathogenic potential of serotype 4b strains. It is likely that the methylase strains in this study are slightly different from 4b strains and may be possible variants. Cytosine methylation has been observed in *Salmonella* [67] and may cause Sau3AI endonuclease resistance in *L. monocytogenes* epidemic clone I and some strains of serotype 1/2a [68]. Using sequence data, Cherry [69] acknowledged different methylation patterns and proposed that in *Escherichia coli* and related enteric bacteria, the inner cytosine are methylated by the Dcm enzyme which causes hypermutation. A comprehensive investigation [70] of cytosine and adenine DNA modifications in several bacteria and other forms of life confirmed the dominance of adenine methylation in bacteria. However, the phylum firmicutes, which includes *L. monocytogenes* recorded more DNA cytosine methylation or modification than actinobacteria, bacteroidetes, and proteobacteria.

## 4. Materials and Methods

### 4.1. Strains, media and DNA extraction

A strain collection of 45 *Listeria monocytogenes* strains stored at -80°C and previously characterized with PCR-serotyping in the Food Microbiology and Safety group of the University of Nottingham were resuscitated and used. Beads containing the strains were removed from the freezer and streaked onto Brain Heart Infusion (BHI, Oxoid UK) agar prepared according to the manufacturer’s instructions. The plates were incubated at 30 °C for 48 h and isolates that emerged were sub-cultured to get single colonies. Genomic DNA for the study was prepared as described previously [7]

### 4.2. Screening of isolates for methylase genes

Amplification for genes of interest was performed in a final volume of 50μl containing 1U of Taq polymerase, 10ng DNA template, 0.2mM deoxyribonucleoside triphosphate, 2mM MgCl2 and PCR Buffer. The forward and reverse primers *meth*F and *meth*R primer (Table 3) was added at a concentration of 1mM. PCR reaction was performed with an initial denaturation step at 94°C for 3 mins followed by 35 cycles of 94°C for 0.40 min, 53°C for 1.15 min and 72°C for 1.15 min and one final cycle of 72°C for 7 mins in a thermocycler (Techne 312). The reaction mixture (5ul) was mixed with 3ul of loading buffer, separated on a 2% agarose gel in TAE buffer, and stained with ethidium bromide before viewing UV light. The products were scored against a 100bp ladder, and strains showing the methylase gene were recorded.

**Table 3.**
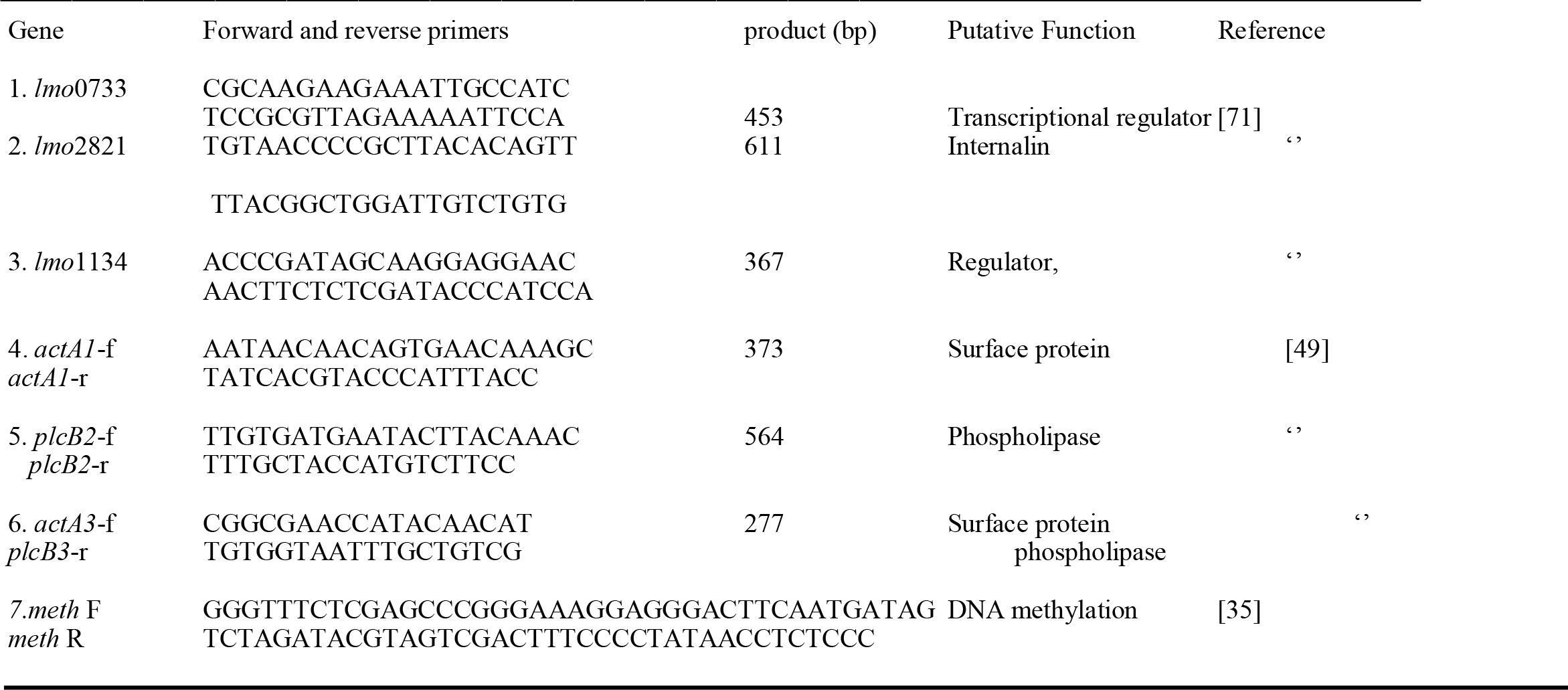
Primers used for this study.

### 4.3. Evolutionary lineage analysis

#### 4.3.1. Lineage classification of *L. monocytogenes* by allelic specific oligonucleotide PCR (ASO-PCR)

The method by Ward et al. [49] was used to perform multiplex ASO-PCR lineage identification based on the virulence gene clusters of *pfrA* genes. Three sets of primers (No 4, 5 and 6 in table 2.6.1) were used. Amplifications were performed in 50μl volumes with 0.5μM concentrations of each primer, 2mM MgCl2, 0.2mM concentrations of each deoxynucleotide triphosphate, 0.5 U of Taq polymerase (Abgene) and 100ng of DNA. Amplifications consisted of 25 cycles of 15s at 94°C, 10s at 56°C and 10 seconds at 72°C following which amplification products were resolved in 1.5% (wt/vol) agarose gel and scored relative to 100bp DNA size ladder.

#### 4.3.2. Differentiation of *L. monocytogenes* serotype 4b under study into evolutionary Lineage I or III

The method of Liu et al. [71] was used to determine if the *L. monocytogenes* 4b strains under study belong to Lineage I or III. PCR primers from virulence-specific genes lmo2821 and lmo1134, species-specific gene lmo0733 and serotype-specific gene ORF 2110 shown in table 2.3 were used. The PCR reaction was done in a reaction volume of 50μl consisting of 0.5U Taq DNA polymerase (Thermo Scientific), 1X PCR buffer 50M dNTPs, 25pmol each primer and 10ng of DNA. Cycling was performed in Techne 312 thermocycler under the following conditions, 94°C for 2 mins once; 94°C for 20 s, 56°C for 20s and 72°C for 45s 30 times and a final extension of 72°C for 2 minutes.

### 4.4. Ribotyping by Southern hybridization

This was carried out as per the manufacturer’s instructions with all kits used.

#### 4.4.1. Restriction enzyme digest

Restriction enzyme digestion with EcoRI (Promega, Madison, U.S.A) was performed by placing 15.3μl of sterile RO water in an Eppendorf. To this 2.5μl of 10x restriction enzyme buffer and 0.2 acetylated bovine serum albumin (BSA) were added, following which 5μl of DNA (approximately 10μg) was added and lastly, 0.5μl restriction enzyme (0.5 units). Samples were mixed and centrifuged for a few seconds to bring contents to the bottom of the tube and incubated overnight at 37°C. The mixture was incubated overnight at 37 °C. Digestion was then assessed on 1.0% agarose in 1x TAE buffer containing ethidium bromide (0.5mgml-1) at 80 vcm^-1^. The ribotype hybridization probe DNA was prepared by reverse transcription of RNA from *Escherichia coli* MRE 600 containing the 16S rRNA operon to get the complementary DNA. Non-radioactive labelling as described in the manufacturer’s kit (Boehringer Manheim) protocol was performed. To a sterile Eppendorf, the following were added. A 2μl hexanucleotide mixture (10x), 2μl reaction buffer for AMV reverse transcriptase enzyme, and 14.25μl distilled water. The mixture was heated at 68°C for 5 minutes and then cooled to room temperature. The following were then added, 2μl 10x dNTP labelling mixture containing DIG-dNTP and 1.5μl (20U/ml) of the reverse transcriptase enzyme. After a quick spin of 30 s, the mixture was incubated at 42 °C overnight and then added to 10 ml DIG-Easy hybridization solution (Roche).

#### 4.4.2. Southern blotting

Blotting of DNA onto a membrane was performed with a Model 785 vacuum blotter (Bio-Rad) following the description of standard transfer in the instruction manual. Restriction digest products (25μl) were electrophoresed through a 0.7% agarose gel at 30V overnight. The gel was photographed, and the distance migrated by each molecular weight marker was noted. The nylon membrane (Hybond N+, Amersham) was placed in sterilized reverse osmosis water at an angle of 45 degrees before wetting with the transfer solution (0.4M NaOH). This was then transferred onto a filter paper which has also been soaked with the transfer solution. The appropriate rubber gasket was then overlaid before the agarose gel was carefully laid over on top. Approximately, 1.5-2.0 litres of transfer solution were poured on top of the gel and then left to blot for 90 min. After blotting, the membrane was washed in 2X SSC for 2 minutes before prehybridization.

Prehybridization prepares the membrane by blocking non-specific nucleic acid binding sites on the membrane. The membrane was placed in a bag containing 20ml of prehybridization solution and then prehybridized at 42 °C for 2 hours following which the hybridization solution containing the DIG-labelled probe was added and hybridized for 20 hr at 42 °C. At the end of the hybridization, the membrane was washed twice, 5 mins per wash in 2X SSC at room temperature to remove the unbound probe and then washed twice, 15 mins per wash in 0.5 X SSC solutions at 68 °C. Colourimetric detection was carried out as described in the manufacturer’s instructions. The colour substrate consisting of 45μl NBT solution and 35μl BCIP solution in 10ml detection buffer was used to incubate the membrane in a sealed plastic bag in the dark until bands were detected. The membrane was photographed and finally washed with water to prevent over-development.

### 4.5. Ancestry of genes associated with cytosine DNA methylation

The complete nucleotide sequence of genes associated with DNA methylation (GenBanknumber AJ302030.1) in *L monocytogenes* was used to search (on 21.12.22) in the BLAST ® suite [72]. The sequence was chosen because it contains cytosine methyl transferase gene *lcmA*. The nucleotide unit consisted of the *lcmA* gene and genes that encodes a heat shock protein, phosphomethyl pyrimidine kinase and thiamine phosphate pyrophosphorylase.

The database was set to nucleotide collection nr/nt and was optimized for highly similar sequences. The algorithm parameters were set to display up to possible 5000 aligned sequences. After the initial shortlisting of 396 sequences from several genera, twenty-four sequences with query coverage of 100 % and over 98% identity were selected for further analysis. These sequences were analysed with the Molecular Evolutionary Genetics Analysis (MEGA) computer software, version 11 [73]. Alignment was built with MUSCLE and a test of phylogeny was carried out using the maximum likely hood statistical method based on the Tamura-Nei model. The bootstrap iterations were set at 1000 iterations after which a phylogenetic tree was constructed to determine evolutionary relationships. The most probable ancestors of *L. monocytogenes* strains harbouring the sequence analysed were identified using the ancestry function. Clonal complex, sequence type, and epidemic clone of some strains were obtained by searching the *Listeria* PasteurMLST database [56].

## 5. Conclusions

This study was carried out to establish the prevalence of strains with methylase genes and their evolutionary history in a set of *L. monocytogenes*. The complete nucleotide sequence of these genes is conserved in *L. monocytogenes* and not yet orthologous in other *Listeria* species and bacteria genera. There may be plasmid-mediated horizontal gene transfer of this methylase sequence in *L. monocytogenes*. Our findings are of epidemiological interest and require broader and continuous surveillance to ascertain if these rare strains are linked with increased pathogenicity, food type or demographic region. In addition, definitive methylation patterns in serotypes and evolutionary lineages will need to be determined.

## Supplementary Materials

The following supporting information can be downloaded at: www.mdpi.com/xxx/s1, Table S1: Blast output with AJ02030.1 Carried out on 21.12.22. (The accession is hyperlinked and can show NCBI details)

## Author Contributions

Conceptualization, O.N and C.R.; methodology, O.N and C.R; software, O.N; formal analysis, O.N and C.R.; investigation O.N and C.R.;.; resources O.N, HO and C.R.; writing—original draft preparation, O.N; writing—review and editing, O.N, HO and C.R All authors have read and agreed to the published version of the manuscript.

## Funding

This research was supported by the DV memorial fund.

## Data Availability

Data sharing is not applicable to this article.

## Conflicts of Interest

The authors declare no conflict of interest.

## Notes

### Competing Interest Statement

The authors have declared no competing interest.

